# Modeling circRNAs expression pattern with integrated sequence and epigenetic features identifies H3K79me2 as regulators for circRNAs expression

**DOI:** 10.1101/392019

**Authors:** Jia-Bin Chen, Shan-Shan Dong, Shi Yao, Yuan-Yuan Duan, Wei-Xin Hu, Hao Chen, Nai-Ning Wang, Ruo-Han Hao, Ming-Rui Guo, Yu-Jie Zhang, Yu Rong, Yi-Xiao Chen, Hlaing Nwe Thynn, Fu-Ling Zhou, Yan Guo, Tie-Lin Yang

## Abstract

Circular RNAs (circRNAs) are an abundant class of noncoding RNAs with widespread, cell/tissue specific pattern. Because of their involvement in the pathogenesis of multiple disease, they are receiving increasing attention. Previous work suggested that epigenetic features might be related to circRNA expression. However, current algorithms for circRNAs prediction neglect these features, leading to constant results across different cells.

Here we built a machine learning framework named CIRCScan, to predict expression status and expression levels of circRNAs in various cell lines based on sequence and epigenetic features. Both expression status and expression levels can be accurately predicted by different groups of features. For expression status, the top features were similar in different cells. However, the top features for predicting expression levels were different in different cells. Noteworthy, the importance of H3K79me2 ranked high in predicting both circRNAs expression status and levels across different cells, indicating its important role in regulating circRNAs expression. Further validation experiment in K562 confirmed that knock down of H3K79me2 did result in reduction of circRNA production.

Our study offers new insights into the regulation of circRNAs by incorporating epigenetic features in prediction models in different cellular contexts.

## Introduction

Circular RNAs (circRNAs) are an abundant class of noncoding RNAs with the widespread, cell-type and tissue specific expression pattern (Memczak et al. 2013; Salzman et al. 2013; Xia et al. 2016). The single-stranded closed circular RNA molecules were first observed in viroid (Sanger et al. 1976). Later, researchers found the circular shape of small RNA structural variants in the cytoplasm of eukaryotic cell with low expression level (Hsu and Coca-Prados 1979), which was considered to be the products of mis-splicing (Cocquerelle et al. 1993). Recent studies have identified a number of exon circularization events across a diversity of species (Cocquerelle et al. 1992; Capel et al. 1993; Salzman et al. 2012), especially in human (Memczak et al. 2013; Salzman et al. 2013; Jeck and Sharpless 2014). The well-known biological function of this class of noncoding RNA is to act as epigenetic regulators that regulate gene expression without changing DNA sequences. Their predominant effect on target gene expression is resulted from their competitive binding affinity with miRNA, which is recognized as miRNA sponges (Hansen et al. 2013a; Hansen et al. 2013b; Guarnerio et al. 2016). Since their significant functions, circRNAs have been reported to be broadly involved in biological processes. And circRNA dysfunction could lead to various diseases, especially cancers (Han et al. 2017; Hsiao et al. 2017; Xia et al. 2018). To date, millions of circRNAs have been identified to be cancer-specific in various cancer cell lines (Zhou et al. 2018).

Analyses of intron sequences flanking circularized exons show enrichment of repeat sequences, which are believed to be necessary for circularization (Dubin et al. 1995; Zhang et al. 2014). Specifically, *Alu* repeats were found to be enriched in the flanking intron regions of circRNA-forming exons with high conservation and were correlated with the human circRNAs formation. The competition between these inverted repeated *Alu* pairs can promote and regulate alternative circularization, resulting in multiple circular RNA transcripts derived from one gene (Jeck et al. 2013; Liang and Wilusz 2014; Zhang et al. 2014). According to the “exon skipping” (Vicens and Westhof 2014; Barrett et al. 2015; Starke et al. 2015) model of exon circularization, a method (Ivanov et al. 2015) was developed to predict circRNAs according to sequence features in the intron region. Besides, some tools also tried to distinguish circRNAs from other lncRNAs based on conformational and conservation features, sequence compositions, *Alu,* SNP densities, and thermodynamic dinucleotide properties (Pan and Xiong 2015; Liu et al. 2016). However, these methods based on genomic sequence features generate indiscriminate predicted results across different tissue/types, which is unable to find the tissue/cell type circRNAs. More importantly, these tools only focus on indicating the circRNAs expression status, and are not capable of predicting circRNAs expression values.

Several studies have used epigenetic or chromatin features to predict gene expression, for example, Karlić *et al.* applied a linear regression model using histone modifications to predict gene expression on human T-cell (Karlic et al. 2010), Dong *et al.* applied a Random Forest Classifier on histone modifications to model gene expression in several human cell lines (Dong et al. 2012), Ritambhara *et al.* constructed a deep-learning model on 56 Roadmap Epigenome Project (RMEC) cell lines to predict gene expression from histone modifications (Singh et al. 2016). Moreover, *Alu* repeats have been reported to be associated with epigenetic features. For example, histone H3 Lysine 9 Methylation (H3-K9) marks were found to be enriched at *Alu* repeats regions in Human Cells (Kondo and Issa 2003; Kondo et al. 2004). *Alu* elements was also found to be bound by two well-phased nucleosomes that contain histones marks of active chromatin, and showed tissue-specific enrichment for the enhancer mark H3K4me1 (Su et al. 2014). Therefore, we hypothesized that the expression of circRNAs could be regulated by epigenetic elements as other noncoding RNAs or protein-coding genes. And interpretation of epigenetic features for known circRNAs could be used for modeling circRNAs expressions and exploring new circRNAs.

In this study, we developed a two-phase pipeline named CIRCScan to model circRNAs expression status and levels based on sequence information and epigenetic features, including *Alu* elements, histone modifications and DNase I hypersensitivity sites (DHSs). We firstly modeled the expression status of circRNAs in GM12878, H1-hESC, HeLa-S3, HepG2, K562 and NHEK cell lines. The predicted accuracy was high with AUC values of 0.78~0.82 for six cell lines. Predicted expressed circRNAs in HeLa-S3 and K562 cell lines were further validated by RNA-seq data. Regression model was then applied to predict circRNAs expression values in different cell lines. The root-mean-square error (RMSE) of models were low with the values of 1.32~1.61 among 6 cell lines and the predicted circRNAs expression values also showed correlation with the actual expression values with the Pearson’s correlation coefficient *r* (PCC) of 0.38~0.45. Among all features used, K3K79me2 showed high importance in modeling both circRNAs expression status and levels across all cell lines. Knocking down of H3K79me2 in K562 cell line led to a significant reduction of circRNAs numbers and expression levels, indicating the important role of H3K79me2 in regulating both circRNAs expression status and levels. Our results demonstrated that the combination of sequence and epigenetic features could be used to model circRNAs expression and explore novel circRNAs in various cell lines.

## Results

To accurately predict circRNAs expression status and levels with the combination of complex factors, we applied a two-phase machine learning pipeline (Figure 1).

**Figure 1.**
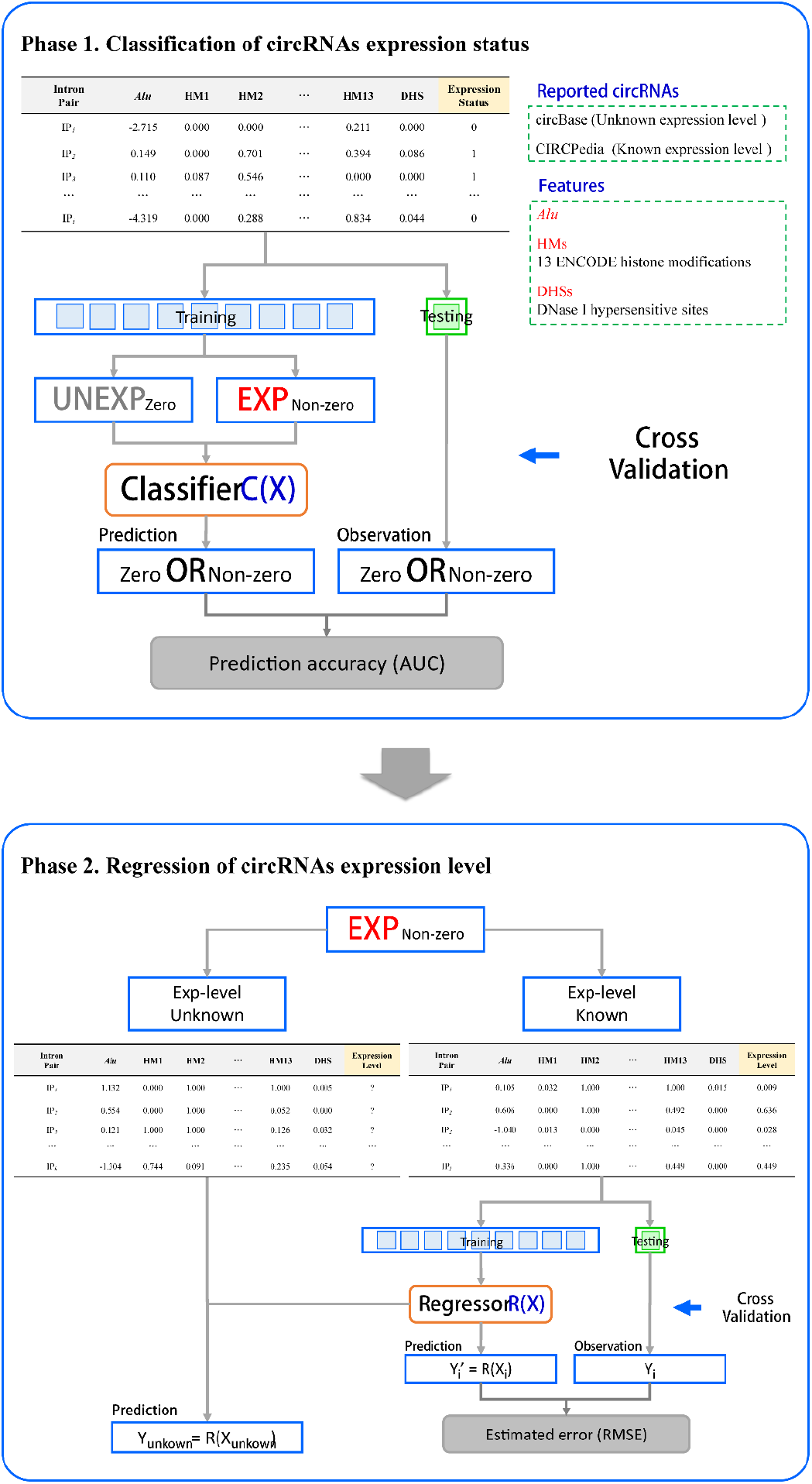
Two-phase machine learning pipeline of circRNAs expression prediction. *Alu* sequence feature and epigenetic features including 13 histone modifications and DHSs were used for model construction. For the first phase, classification algorithm was applied to model the expression status of circRNAs expression. Data of reported known circRNAs were downloaded from circBase and CIRCPedia databases. CIRCPedia also provides the circRNAs expression status. 10-fold cross-validation was carried out to reduce the biases during modeling. All training data labeled as expressed (non-zero) and unexpressed (zero) were randomly partitioned into 10 equal size subsets, of which 9 subsets were used to train model and one was retained for testing. A classifier was then trained to distinguish expressed circRNAs (FIPs) from all circRNAs (FIPs) with training set and tested in testing set. AUC score was used as the index to evaluate model performance. The model was then used to predict expressed circRNAs from whole genome. In the second phase, a regressor was trained to model and predict circRNAs expression values. For those expressed circRNAs in CIRCPedia with known expression levels, we applied regression algorithm on these training data to model circRNAs expression levels. 10-fold cross-validation was also used when constructing model. Model performance was evaluated by the estimated error (RMSE), and the final model was used to predict the expression levels of those (predicted) expressed circRNAs. DHSs: DNase I hypersensitive sites; FIPs: flanking intron pairs; AUC: area under ROC curve; RMSE: area under ROC curve. The process of feature selection for model optimization is not shown in the figure.

### Acquisition and annotation of labeled circRNAs intron pairs

For preparing machine learning data sets within both classification and regression modeling phases, we performed a two steps’ screening of introns and intron pairs intervals’ lengths. Eventually, we obtained 1,508,268 intron pairs in total.

After mining circBase (Glazar et al. 2014) and CIRCpedia (Zhang et al. 2016) database, we obtained 21,883, 18,533, 22,053, 16,007, 31,775 and 13,451 labeled positive flanking intron pairs (FIPs) for classification models in GM12878, H1-hESC, HeLa-S3, HepG2, K562 and NHEK cell lines, respectively. Considering the intron length and distance of introns (pairs) to promoters, labeled negative FIPs were obtained by the strategy of stratified sampling as the same number as positive FIPs. Totally, there were 43,766, 37,066, 44,106, 32,014, 18,120, 63,550, and 26,902 intron pairs applied for the model training and testing. For the phase of modeling circRNAs expression levels, we referred to the mapped back-splice junction Reads Per Million mapped reads (RPM) values in CIRCpedia database to evaluate the circRNAs expression levels. 13037, 10431, 16156, 8967, 17996 and 7571 circRNA FIPs with expression levels were selected from all intron pairs for each cell line in GM12878, H1-hESC, HeLa-S3, HepG2, K562 and NHEK, respectively.

A total of 15 genomic and epigenetic features were used to annotate all intron pairs in 6 cell lines, which included *Alu* elements, 13 histone modifications and DHSs for GM12878, H1-hESC, HeLa-S3, HepG2, K562 and NHEK (see details in Table 1).

**Table 1.** Features used for model training in GM12878, H1-hESC, HeLa-S3, HepG2, K562 and NHEK.

| Feature Type (number) | Features |
| --- | --- |
| Sequence (1) | *Alu* |
| Histone modifications (13) | CTCF |
|  | EZH2_(39875) |
|  | H2A.Z |
|  | H3K27ac |
|  | H3K27me3 |
|  | H3K36me3 |
|  | H3K4me1 |
|  | H3K4me2 |
|  | H3K4me3 |
|  | H3K79me2 |
|  | H3K9ac |
|  | H3K9me3 |
|  | H4K20me1 |
| Open chromatin (1) | DNaseI hypersensitive sites (DHSs) |

### Prediction of circRNAs expression status

#### Predicted expression status of circRNAs with high accuracy in different cell lines

In the first phase, we constructed classification models to determine the circRNAs’ expression status. The circRNAs expression status were labeled as expressed (ON) and not expressed (OFF). Among all tested models, we found that random forest (rf) performed significantly superior to others with higher AUC of 0.7981, 0.7779, 0.7876, 0.7820, and 0.7994 for GM12878, H1-hESC, HeLa-S3, HepG2, and NHEK in feature selection (Figure 2A). The model of K562 showed the best performance with AUC of 0.8223. Thus, we chose rf as the final method. Notably, all these 6 cell lines showed a similar pattern. That is, when no more than 3 features were used, the AUC of rf increased significantly along with the number of features increasing. However, the AUC barely changed when more than 3 features were used. Model performances are shown in Table 2. We then ranked all features’ importance in each model, and result showed that, *Alu,* H3K36me3 and H3K79me2 were the top 3 features consistently in all 6 cell lines (Figure 2B), which suggested that these 3 features might play important roles on the process of circRNAs formation. In addition, to explore whether these features were predictive for all cell lines indiscriminatingly or there were different regulation patterns among different cell lines, we performed the cross prediction. Model was trained in one cell line and then was used to predict circRNAs in other cell lines. The results showed that the performance of cross prediction was comparable to which were trained in the original cell line (Figure 2C). And it also indicated that the features were indiscriminatingly predictive and the model could be generally used for predicting circRNAs expression in different cell lines.

**Figure 2.**
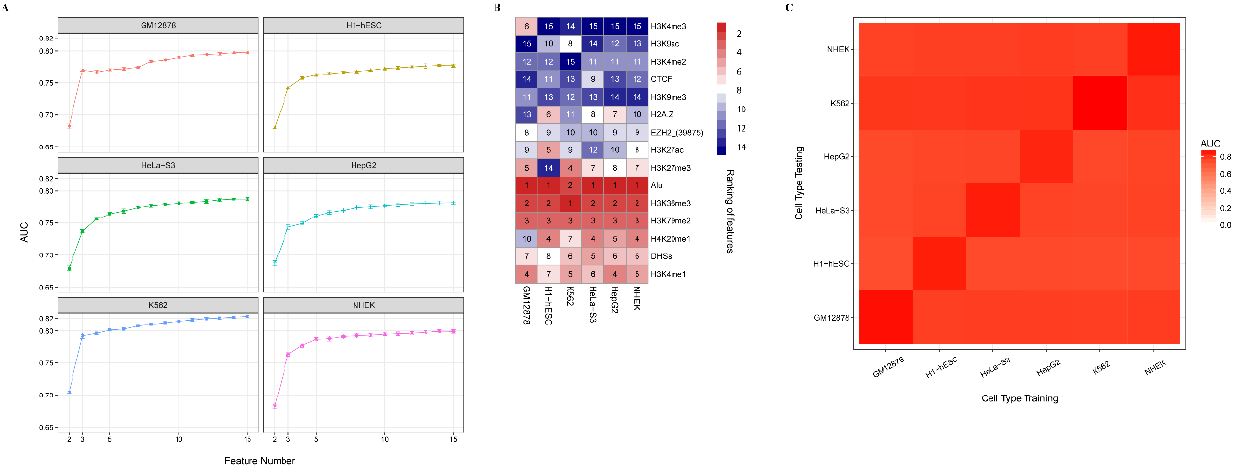
(A) Performance (AUC) and feature selection of rf classification models in GM12878, H1-hESC, HeLa-S3, HepG2, K562 and NHEK. Feature selection was used to find optimal combination or subset of features with the best performance. (B) Predictive importance for features of rf classification models in GM12878, H1-hESC, HeLa-S3, HepG2, K562 and NHEK. Feature importance range from 1 to 15 with the colors range from dark blue to dark red. (Ranking of each feature is shown in corresponding box). (C) The predicted accuracy (AUC) of cross-prediction among GM12878, H1-hESC, HeLa-S3, HepG2, K562 and NHEK for classification models. AUC values decreased with color changed from red to white.

**Table 2.** Summary statistics for the classification models (rf) of GM12878, H1-hESC, HeLa-S3, HepG2, K562 and NHEK.

| Cell Line | GM12878 | H1-hESC | HeLa-S3 | HepG2 | K562 | NHEK |
| --- | --- | --- | --- | --- | --- | --- |
| AUC | 0.7981 | 0.7779 | 0.7876 | 0.7820 | 0.8234 | 0.7994 |
| Accuracy | 0.7350 | 0.7204 | 0.7295 | 0.7190 | 0.7619 | 0.7431 |
| Sensitivity | 0.8006 | 0.7369 | 0.7758 | 0.7662 | 0.8103 | 0.7908 |
| Specificity | 0.6694 | 0.7040 | 0.6833 | 0.6717 | 0.7135 | 0.6955 |

AUC: area under ROC curve.

#### Prediction of circRNAs expression status showed significant cell-specificity

For GM12878, H1-hESC, HeLa-S3, HepG2, K562 and NHEK cell lines, we predicted circRNAs using optimized models with selected features in unlabeled set. 361,672, 399,386, 379,705, 372,310, 327,137, and 413,686 intron pairs were predicted to flank expressed circular exons that could promote exons circularization in unlabeled set of 6 cell lines respectively (Table S1). Totally, there were 735,904 nonredundant predicted expressed circRNAs (FIPs) in all 6 cell lines. Among those, 30.8% (226,343) circRNAs were predicted to be expressed in only one cell line which were considered as cell-specific, while only 13.8% (102,165) were predicted to be expressed across 6 cell lines. This result showed significant cell-specificity of predicted circRNAs in different cell lines.

#### Validating predicted expression status with RNA-seq data

To illustrate the reliability of our prediction, we further performed the RNA-seq analysis of 6 human acute myelocytic leukemia (AML) samples and HeLa cell, to validate the predicted circRNAs expression status in K562 and HeLa cell lines. We detected 9,834 and 1,853 circRNAs in K562 and HeLa-S3 cell lines with relative high abundance, 80.30% (7,897) and 80.52% (1,492) of which were predicted as expressed circRNAs in K562 and HeLa-S3, respectively. We predicted a big majority of authentically expressed circRNAs, which indicated the high accuracy and reliability of our method.

### Modeling of circRNAs expression levels

#### CircRNAs expression levels could be well modeled by quantitative models

Regression algorithms were then applied to model and predict circRNAs expression levels. For each cell line, we estimated error (RMSE) and the correlation (PCC) between observed and predicted expression values to evaluate the model performance. Random forest (rf) showed the lowest RMSE and highest PCC values in all 6 cell lines when as many of features were included in our models (Figure 3A). RMSE values ranged from 1.33 (H1-hESC) to 1.61 (HeLa-S3) and PCC were from 0.38 to 0.45 (Figure 4, Table 3). H1-hESC showed the highest correlation with the lowest RMSE and highest PCC value. We evaluated the feature importance in each cell line. As shown in Figure 3B, H3K79me2 was among the top three important features in all 6 cell lines, suggesting the universal function of H3K79me2 in modulating circRNAs expression levels in different cell lines. We also performed the cross-prediction on circRNAs expression levels, and there was a significant decrease of predicted correlation of other cell lines compared with the original cell line (Figure 3C). Compared with classification models, most features of models showed different predicting power and were superior in predicting circRNAs expression levels to other features in different cell lines, suggesting different modulation patterns of circRNAs expression levels exist among different cell lines.

**Figure 3.**
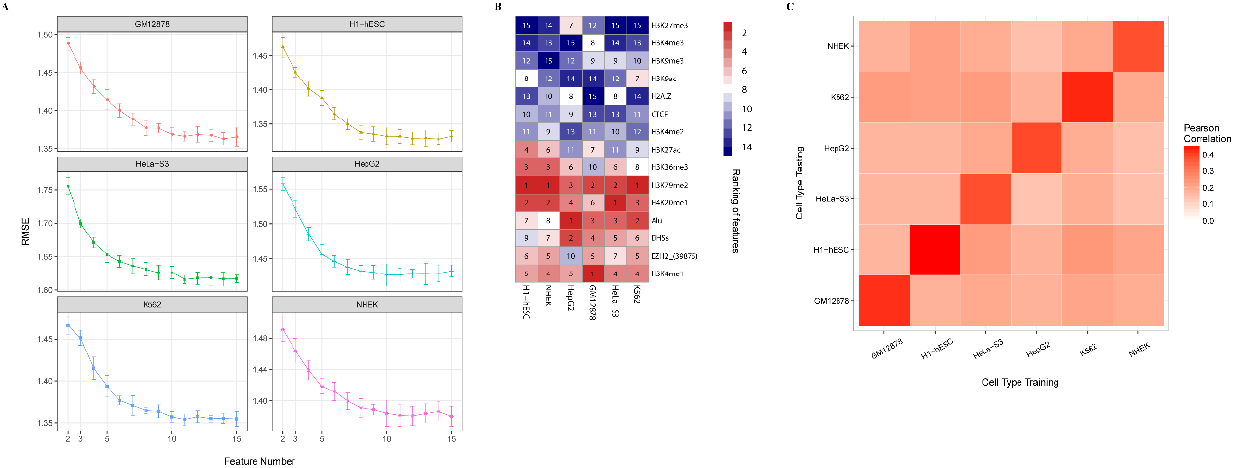
**(A)** Performance (RMSE) and feature selection of rf regression models in GM12878, H1-hESC, HeLa-S3, HepG2, K562 and NHEK. Feature selection was used to call for optimal combination or subset of features with the best performance. **(B)** Predictive importance for features of rf regression models in GM12878, H1-hESC, HeLa-S3, HepG2, K562 and NHEK. Feature importance range from 1 to 15 with the colors range from dark blue to dark red. (Ranking of each feature is shown in corresponding box). **(C)** The predicted accuracy (RMSE) of cross-prediction among GM12878, H1-hESC, HeLa-S3, HepG2, K562 and NHEK for regression models. RMSE values decreased with color changed from red to white.

**Figure 4.**
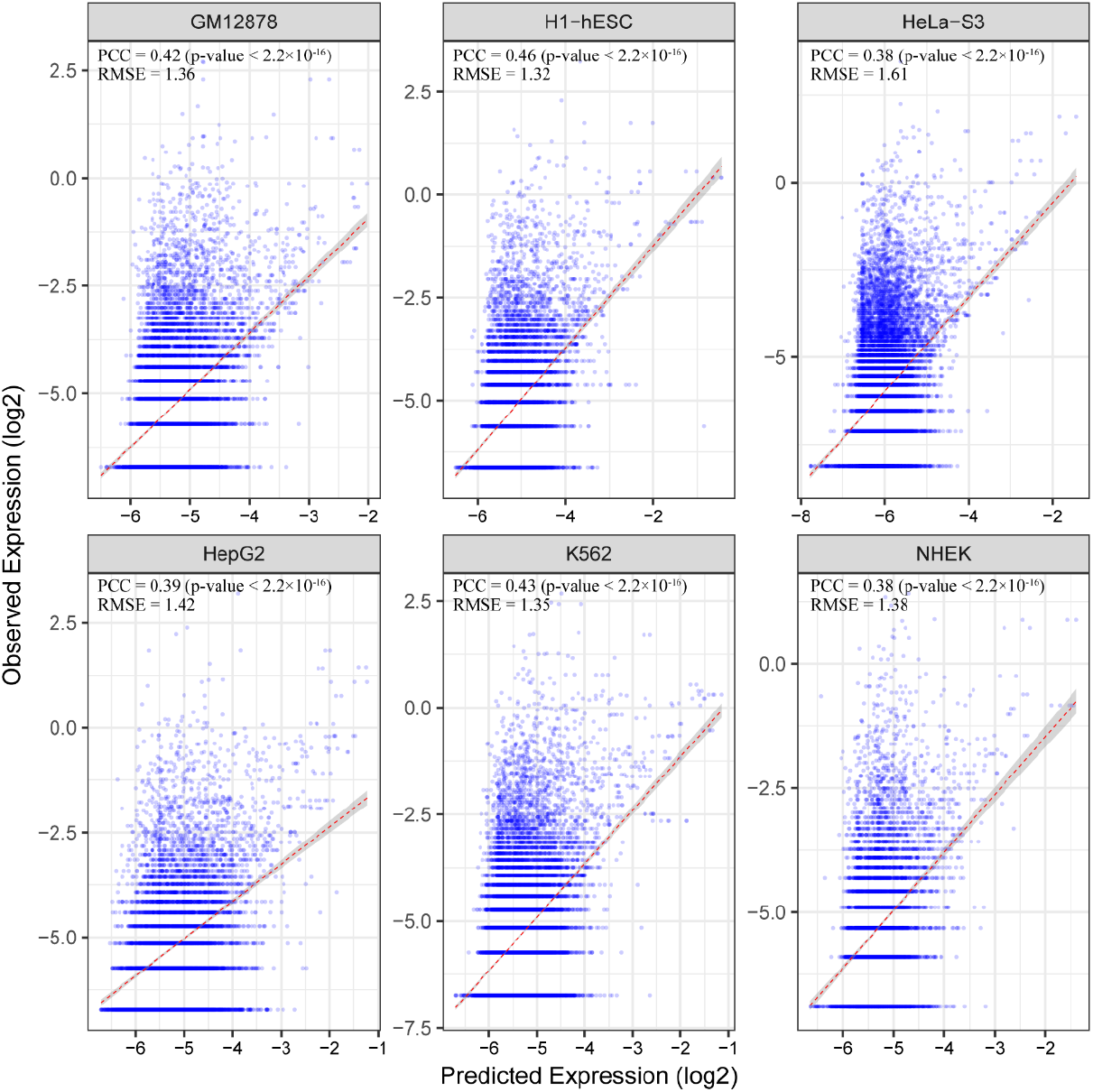
Predicted and observed expression levels (log2 transformed) of circRNAs in GM12878, H1-hESC, HeLa-S3, HepG2, K562 and NHEK. RMSE: root mean square error; PCC: Pearson correlation coefficient. (p-value < 2.2e-16)

**Table 3.** Summary statistics for the regression models (rf) of GM12878, H1-hESC, HeLa-S3, HepG2, K562 and NHEK.

| Cell Line | GM12878 | H1-hESC | HeLa-S3 | HepG2 | K562 | NHEK |
| --- | --- | --- | --- | --- | --- | --- |
| RMSE | 1.36 | 1.33 | 1.61 | 1.42 | 1.35 | 1.38 |
| PCC | 0.42 | 0.45 | 0.38 | 0.39 | 0.43 | 0.38 |

RMSE: root mean square error; PCC: Pearson correlation coefficient.

#### Validating predicted expression levels with known circRNAs in circBase

To demonstrate the predicted accuracy of our circRNAs expression levels, we firstly mined circBase database to verify the circRNAs expression levels. We compared the predicted expression levels of circRNAs in circBase with those were unreported in the data set to see whether there were significant differences between these two groups in predicting expression levels. It showed that the predicted expression levels of circRNAs in circBase were higher than those in unreported group significantly (p-value < 2.2e-16) in all 6 cell lines (Figure 5). We further predicted the expression levels of the circRNAs that were predicted to be expressed and non-expressed. The results showed a higher expression of the predicted expressed circRNAs (Figure S1). Taken together, these evidences indicated that our method of predicting circRNAs expression levels can reflected the authenticable circRNAs expression levels.

**Figure 5.**
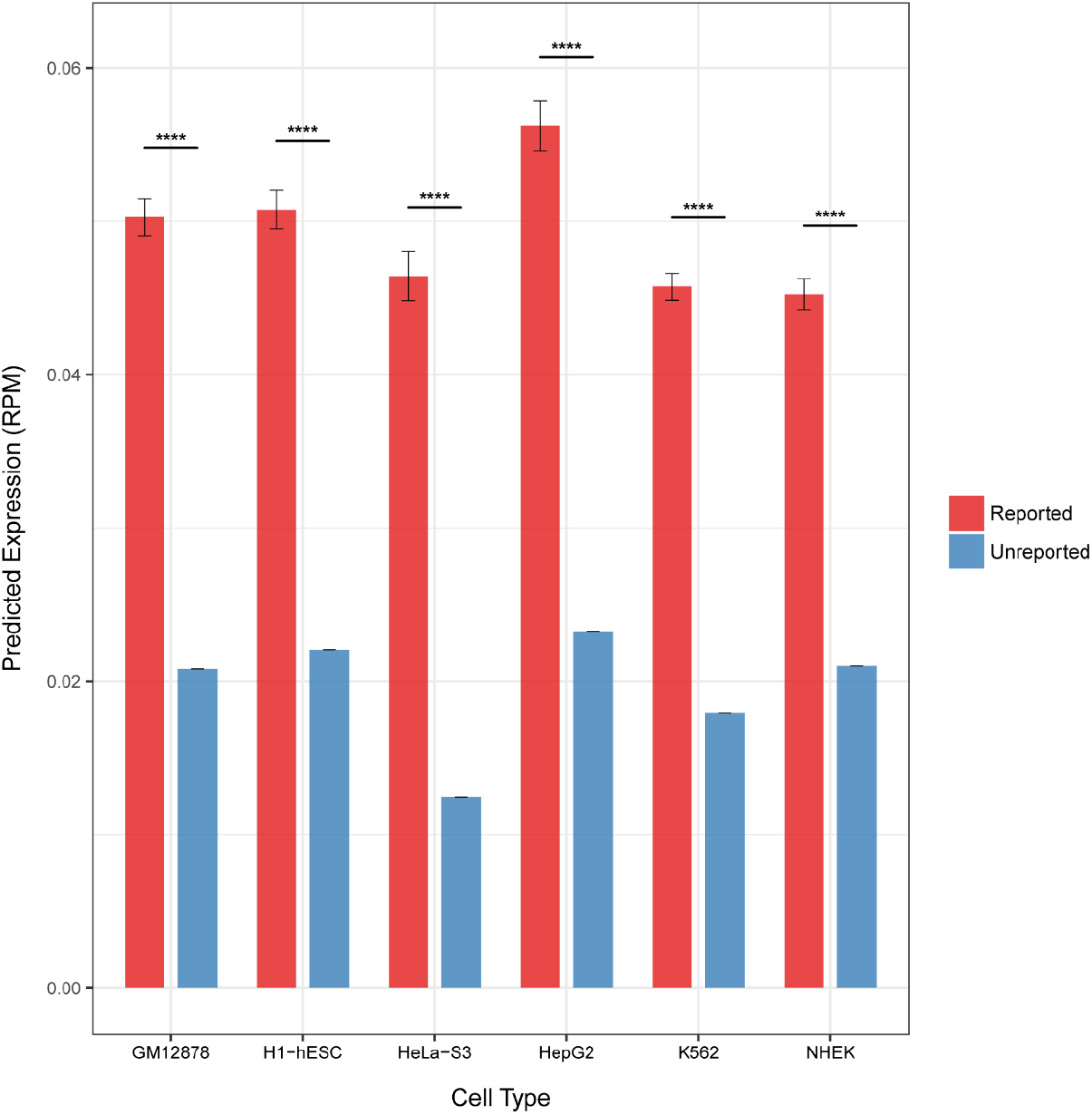
Predicted expression levels of reported and unreported circRNAs in GM12878, H1-hESC, HeLa-S3, HepG2, K562 and NHEK. Reported circRNAs: circRNAs reported in circBase without expression level data. (p-value < 2.2e-16)

### Comparing prediction accuracy of classification and regression models trained with all features, epigenetic features, and only *Alu* elements

To further explore the predictive power of epigenetic features in distinguishing expressed circRNAs from other regions and modeling circRNAs expression levels, we compared the prediction accuracy of models by feature groups of all features, epigenetic features excluding *Alu, Alu* only, and to find whether the using of epigenetic features can significantly improve model performance. Epigenetic features showed superior prediction power than only using *Alu* when modeling circRNAs expression status, and the AUC of epigenetic models was significantly higher than model with only *Alu.* There was also a significant decrease of AUC after removing epigenetic features in all cell lines (Figure S2A). For predicting circRNAs expression levels, the performance of models with epigenetic features showed lower RMSE values, which was not different with models using all features. However, models trained with only *Alu* showed significant decrease of predicted accuracy (higher RMSE values) than models trained with all features or epigenetic features in all 6 cell lines (Figure S2B). These results indicated the importance of epigenetic features in modeling both circRNAs expression status and expression levels, as well as the superiority of epigenetic features than *Alu* in predicting circRNAs expression. Moreover, the addition of epigenetic features in models significantly improved the predictive power.

### CircRNAs expression could be repressed by knocking down the H3K79me2 mark

To validate the functional effect of H3K79me2 on regulating circRNAs expression, we conducted a knock-down experiment of H3K79me2 mark in K562 cell line. CircRNA candidates detected in RNA-seq data were filtered by reads count (> 1 reads) and H3K79me2 annotation (IP_score_*H3K79me2*_ > 0) to make it comparable between control and H3K79me2 knock-down groups. Compared with control group, there was a significant reduction of circRNAs numbers and expression levels after H3K79me2 knocking down. Among 519 circRNAs with H3K79me2 mark in flanking intron pairs in control group, 294 (56.6%) circRNAs were failed to be detected in the knock-down cell line. For the rest of 225 circRNAs, 7 circRNAs were differential expressed (p-value < 0.05), and 5 of them were down regulated in knock-down group. Overall, 299 circRNAs showed decreased expression after H3K79me2 knock-down treatment, accounting for a percentage of nearly 60% circRNAs in untreated K562 cells. This result supported our finding that the H3K79me2 is functionally associated with circRNAs expression regulation, demonstrating that the circRNAs expression was predictable by a set of epigenetic features.

### Comparison with other tools for predicting circRNAs

Compared with previous published tools or methods (Table 4), CIRCScan is more applicable for predicting circRNAs in the following aspects: firstly, CIRCScan is capable of modeling and predicting both circRNAs expression status and expression levels with high accuracy, while other tools used alignment or classification algorithms could only predict the expression status. Secondly, epigenetic and sequence features were both included in our model, which is able to account for the cell specificity. In comparison, other tools only using genomic information could only generate non-cell-specific results. Thirdly, CIRCScan predicted circRNAs expression with the consideration of alternative splicing and different circular isoforms derived from one gene. Moreover, some tools were restricted to only distinguish circRNAs from lncRNAs or constitutive exons, while CIRCScan could be widely used to predicted circRNAs in genome-wide.

**Table 4.** Comparison of previous studies (tools and methods) for the task of predicting circRNAs without RNA-seq data.

| Tool name | Strategy | Features | Feature number | Best model | Expression level | Cell specificity | Alternative back-splicing | Predictive range | Ref |
| --- | --- | --- | --- | --- | --- | --- | --- | --- | --- |
| Ivanov et al. | Alignment | RCMs | 1 | — | × | × | ✓ | Whole genome | (Ivanov et al. 2015) |
| PredcircRNA | ML | Sequence features | 188 | RF | × | × | × | LncRNAs | (Pan and Xiong 2015) |
| predicircrnatool | ML | Conformational Thermodynamic | 125 | SVM | × | × | × | Constitutive exons | (Liu et al. 2016) |
| CIRCScan | ML | Sequence feature<br>Epigenetic features | 15 | RF | ✓ | ✓ | ✓ | Whole genome | — |

RCMs: reverse complementary matches; WG: whole genome; ML: machine learning; RF: Random Forest; LncRNAs: long noncoding RNAs; SVM: Support vector machine; CE: constitutive exons.

## Discussion

Our study placed emphasis on the effect of sequence and epigenetic features in intron regions of human genome on regulating circRNAs expression and attempted to construct a new method to predict circRNAs expression in different cell lines, and moreover, to explore new regulators that are important for circRNAs expression modulation.

In this study, we introduced classification and regression algorithms in our pipeline and successfully predicted both circRNAs expression status and expression levels in different cell lines with high accuracy. Notably, we observed significant difference between these two layers of data. The selected features of regulating expression status was almost the same among different cell lines, while the features playing essential role in expression levels regulation varied a lot across cell types. Compared with previous study that modeled the expression of gene linear isoforms using chromatin features (Dong et al. 2012), we reported discrepant patterns of regulating expression of these two types of gene transcriptional products. It was found that H3K9ac and H3K4me3 showed the highest importance in identifying liner RNAs expression status across multiple cell lines while H3K79me2 and H3K36me3 were more important for regression of expression levels. In comparison, we found that *Alu,* H3K36me3 and H3K79me2 histone marks played the most important role in modeling circRNAs expression status, while the expression levels could be predicted by different groups of features. The results showed that H3K79me2 was important for modeling both circRNAs expression status and levels in various of cell lines. The different feature importance between modeling linear and circular isoforms indicates that there could be different transcriptional regulatory mechanisms between linear and circular isoforms which could be explained by disparate splicing process.

*Alu* elements have already been reported to affect gene conversion and alterations in gene expression (Batzer and Deininger 2002), and are highly correlated with the formation of human circular RNAs (Jeck et al. 2013). It was also confirmed by our results that *Alu* showed the most importance in predicting circRNAs expression status in almost all cell lines. While it was less important for modeling circRNAs expression levels. In this study, we added epigenetic features in modeling circRNAs expression status and levels, and there was a significant improve of model performance especially for the expression levels. Actually, epigenetic features made even higher contribution compared with *Alu* and genomic features in modeling both circRNAs expression status and expression levels (Figure S2). All these results confirmed that the combining sequence and epigenetic features could better model circRNAs expression in various cell lines.

Histone mark of H3K79me2 showed high importance in modeling both circRNAs expressions status and levels, indicating the virtual role of epigenetic elements in transcriptional regulation. Characterization of chromatin states in human genome showed that, H3K79me2 marked transcription-associated states strongly enriched for spliced exons and *Alu* repeats (Ernst and Kellis 2010). On one hand, H3K79me2 was reported to be positively associated with transcriptionally active genes (Onder et al. 2012). As a mark of the transcriptional transition and elongation region, H3K79me2 could influence alternative splice site choice by modulating RNA polymerase II (PolII) elongation rate (Guenther et al. 2007). H3K79me2 marks enriched in genes with high elongation rates, and the high elongation rates had a positive correlation with distance from the nearest active transcription unit, and low complexity DNA sequence such as *Alu* of genome short interspersed element (SINE) (Veloso et al. 2014). On the other hand, inhibition or slowing of canonical pre-mRNA processing events has been proved to shifts the normal protein-coding genes products toward circular RNAs isoforms and increased the circRNAs output (Liang et al. 2017). Therefore, the absence of H3K79me2 may not only influence the pre-mRNA splicing but all transcriptional regulation process.

Our study implemented a two-phase machine learning strategy to characterize the FIPs of exon(s) regions with sequence and epigenetic features, and finally accurately predict circRNAs expression in 6 ENCODE cell lines. We identified H3K79me2 as a new epigenetic regulator for circRNAs expression modulation, which was further validated by the H3K79me2 knockdown experiment in K562 cell. Collectively, our work provides a new strategy for modeling circRNAs expression pattern in various cell lines, and offers new insights on the mechanisms of circRNAs expression regulation.

## Methods

### Acquisition of known circRNAs and screening for genomic intron pairs

The data sets of known circRNAs of different cell lines were downloaded from circBase (http://www.circbase.org) (Glazar et al. 2014) and CIRCpedia (http://www.picb.ac.cn/rnomics/circpedia) (Zhang et al. 2016) databases. CircRNAs from circBase and CIRCpedia circRNAs were mapped with all intron pair intervals using Bedtools (Quinlan and Hall 2010) (Bedtools 2.25.0 with parameters: intersect –f 1.0 –F 1.0) and the acquired circRNAs were used as training set for feature generation. All reference exons and introns were extracted from UCSC gene annotation of Human genome hg19 assembly with custom python scripts. Introns with genomic length over 300 bp (according to the length of *Alu* repeats) were retained. Introns in a common transcript were paired, and duplicate intron pairs of same position in different transcripts were removed. For these intron pairs, we removed those with interval length of two introns shorter than 50 bp or longer than 2M bp according to the genomic length of known circRNAs from circBase and CIRCpedia.

### Feature annotation

*Alu* elements data (RepeatMasker) of Human Genome hg19 were downloaded from USCS Table Browser (http://genome.ucsc.edu/cgi-bin/hgTables). Functional epigenomic data of 6 human cell lines were downloaded from Encyclopedia of DNA Elements (ENCODE) Project Consortium (Consortium 2012) (ftp://hgdownload.cse.ucsc.edu/goldenPath/hg19/encodeDCC/), including GM12878, H1-hESC, HeLa-S3, HepG2, K562 and NHEK. Features were: *Alu* repeats, histone modifications, and DHSs elements.

Bedtools (Bedtools 2.25.0) “intersect” sub-command was used to overlap the *Alu* repeat regions with genomic intron regions (parameters: bedtools intersect –s/-S -F 1.00 -a -b -wa -wb). For *Alu* repeats, we annotated the intron pairs with IP_score_*Alu*_ by considering the competition of inverted repeated complementary sequences (IR*Alus*) within (IRAlus_*within*_) or across flanking introns (IR*Alus*_*across*_) and the genomic length of introns. IP_scoreA_*Alu*_ was defined as:

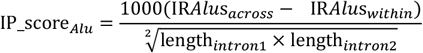

IR*Alus*_*across*_ – IR*Alu*s_*within*_: difference of IR*Alu*s_*acmss*_ axnd IR*Alu*s_*within*_ number

Regions of each selected ENCODE epigenetic element in 6 cell lines were firstly extracted and then the overlapped BED/GFF/VCF entries were merged into a single interval using Bedtools “merge” sub-comm. Merged regions were then intersected with intron regions (parameters: bedtools intersect -a -b -wao). For each of the epigenetic feature, we annotated the intron pairs by IP_score_*epi*_ of which the density of features in intron regions was take into consideration. IP_score_*epi*_ was defined as:

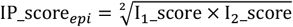

And I_*i*__score_*epi*_ (score of the *i* intron within a intron pair) was defined as:

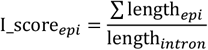

Σ length_*epi*_ referred to the summation of non-overlapping length of an epigenetic feature in an intron region. After annotating intron pairs with each feature, all annotation of sequence and epigenetic features were combined together eventually.

### Training data preparation

In classification models, all intron pairs (intervals) were firstly divided into expressed circRNAs FIPs (positive sets) and unknown intron pairs by comparing their genomic locations with known circRNAs FIPs in circBase (Glazar et al. 2014) and CIRCpedia (Zhang et al. 2016) database mentioned before. When preparing positive and negative FIPs, introns length and distance of introns (pairs) to promoters were taking into consideration to avoid potential bias. For intron length, we used stratified sampling and set several length ranges (parts) and then assigned intron pairs into each part according to the sum of introns length. P-values of the flanking introns length for positive and negative sets were then calculated to evaluate the results of stratified sampling (Table S2). For distance of introns (pairs) to promoters, we divided the FIPs into two parts according to whether they had proximal promoters (within 1M bp region) or not to make sure the proportions of intron pairs with proximal promoters in positive and negative sets were the same. Promotor region was defined as the +/− 1K bp region of transcription start site (TSS). Randomly sampled negative sets were then combined with positive sets as the training data sets for machine learning. In regression models, all intron pairs were divided into two groups according to whether there were expression values available. Those (circRNA) FIPs with expression levels were used to constructed regression models and then applied to predict expression values of the predicted expressed circRNAs.

### Model generation, evaluation and optimization

Five different types of widely used classification algorithms of machine learning, including linear discriminant analysis (lda), naive bayes (nb), neural network (nnet), random forest (rf) and bagged CART (treebag) were applied in constructing classification models. In phase II, algorithms of nnet, rf and treebag were further used to construct regression models. All these algorithms were implemented in the “caret” package in R (version 3.2.4). To reduce the bias during sampling, 10-fold cross-validation was carried out in all classification and regression models. All training data was randomly partitioned into 10 equal size subsets, of which 9 subsets were used to train model and one retained for testing. Function “varImp” was used to calculate the importance of each feature. For random forest classification models, *MeanDecreaseGini* was used as a measure of variable importance based on the Gini impurity index (Breiman 2001), and permutation-based MSE reduction (“permutation importance”) was used as variable importance metric for random forest regression (Strobl et al. 2008). Parameter tuning was also applied for each algorithm to get the best parameters and optimize the model performance. To remove the redundant information and obtain the optimal subset of features, we used the recursive feature elimination (Guyon 2002). To improve the computational efficiency of model training, we applied the parallel processing frameworks in R (R package “doParallel”) during model generation. Finally, the best performing subset was used to generate the final model. The model was then applied to predict the expression status and levels of circRNAs by the function of “predict” packaged in “caret”.

### Evaluation of model performance

Several generally used indexes were applied in our study to evaluate the performance of models. For classification models, sensitivity (recall), specificity, accuracy, precision and area under the curve of ROC (AUC, sensitivity versus 1 – specificity) were used to evaluate predicted accuracy. True positive (TP) and false negative (FN) represent the numbers of positive instances predicted to be positive or negative separately. Similarly, false negative (FP) and true negative (TN) refer to the numbers of negative instances which are predicted to be positive or negative respectively. AUC score was applied as the core index to evaluate model performance and other indexes for references. Root-mean-square error (RMSE) values and Pearson’s correlation coefficient *r* (PCC) were calculated to evaluate the regression models performance. RMSE refers to:

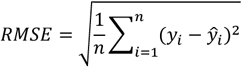

n: number of observations; yi: observed value; ŷi: predicted value.

### Cell culture

Human embryonic kidney cells (HEK293T) were cultured in Dulbecco’s modified Eagle’s medium (DMEM) and Human erythroleukemia cell line K562 cells were cultured in Roswell Park Memorial Institute (RPMI) 1640 (Biological Industries, Israel). All of the cells were cultured containing 10% fetal bovine serum (FBS) (Biological Industries, Israel) with 1% penicillin/streptomycin at 37°C, 5% CO2.

### Construction of shRNA knock-down plasmids

*AF10* gene knock-down shRNA target sequence were reported in NCBI probe database https://www.ncbi.nlm.nih.gov/probe/. *AF10* siRNA target site is 5’-CCTGCTGTTCTAGCACTTCAT-3’ (Deshpande et al. 2014). We used mi30 backbone to expression the shRNA and constructed it into pCDH-CMV-MCS-EF1 α-puro lentivirus plasmid. All sequences of the primers for *AF10* shRNA and mi30 shRNA backbone are shown in the Table S3.

### Transfection and lentivirus

The lentivirus-mediated shRNAs targeting *AF10* was transfected into HEK293T cells with lentivirus package helper plasmids (psPAX2 #12260 and pCMV-VSV-G #8454). After 48h and 72h collected the supernatant medium and kept in -80°C. All plasmids transfected into HEK293T cells were used ViaFect™ transfection reagent (Promega, USA). Nextly, 8×10^5^ K562 cells were infected with the supernatant medium for 72h in 6-well plates and selected by puromycin (1.75μg/ml).

### RNA extraction, quantitative real-time PCR

Total RNA was extracted from K562 cells using Trizol (Life Technologies, USA) with DNase I digestion according to the manufacturer’s instructions. The reverse transcription reaction was performed with HiScript Q RT SuperMix for qPCR (Vazyme Biotech, China) following manufacturer’s instructions in C1000 TouchTM Thermal Cycler (Bio-Rad, USA). Next, the expression of *AF10* mRNA level was detected by SYBR Green PCR kit (QuantiFast, Germany), according to manufacturer’s instruction in CFX ConnectTM Real-Time System (Bio-Rad, USA). The *ACTB* was used as endogenous control. All sequences of the primers used are shown in the Table S3. Results of qPCR are shown in Figure S3.

### RNA quantification and qualification

RNA degradation and contamination was monitored on 1% agarose gels. RNA purity was checked using the NanoPhotometer spectrophotometer (IMPLEN, USA). RNA concentration was measured using Qubit RNA Assay Kit in Qubit 2.0 Flurometer (Life Technologies, USA). RNA integrity was assessed using the RNA Nano 6000 Assay Kit of the Bioanalyzer 2100 system (Agilent Technologies, USA).

### Library preparation and sequencing of human K562 cells

Ribosomal depleted sequencing libraries were generated for all samples using NEBNext UltraTM RNA Library Prep Kit for Illumina (NEB, USA) following manufacturer’s recommendations and index codes were added to attribute sequences to each sample. Fragmentation was carried out using divalent cations under elevated temperature in NEBNext First Strand Synthesis Reaction Buffer (5X). First strand cDNA was synthesized using random hexamer primer and M-MuLV Reverse Transcriptase (RNase H-). Second strand cDNA synthesis was subsequently performed using DNA Polymerase I and RNase H. Remaining overhangs were converted into blunt ends via exonuclease/polymerase activities. After adenylation of 3’ ends of DNA fragments, NEBNext Adaptor with hairpin loop structure were ligated to prepare for hybridization. The library fragments were purified and size-selected for 250-300 bp cDNA fragments with AMPure XP system (Beckman Coulter, Beverly, USA). Then 3 μl USER Enzyme (NEB, USA) was used with size-selected, adaptor-ligated cDNA at 37°C for 15 min followed by 5 min at 95 °C before PCR. PCR was performed with Phusion High-Fidelity DNA polymerase, Universal PCR primers and Index (X) Primer. At last, PCR products were purified (AMPure XP system) and library quality was assessed on the Agilent Bioanalyzer 2100 system. Library preparations were then sequenced using Illumina HiSeq X Ten and 150 bp paired-end reads were generated.

### RNA-seq data preparation for human leukemia samples

RNA-seq data of 6 in-house acute myelocytic leukemia (AML) samples were used to detect circRNAs. Total RNA was extracted from monocytes of 6 acute myeloid leukemia samples using Trizol Reagent (Life Technologies, USA). Total RNA was then depleted of ribosomal RNA using a Ribominus kit (Invitrogen, USA) according to the manufacturer’s instructions. RNase-R treatment was carried out for 1 hour at 37°C using RNase R (Epicenter, USA) 1.5 U/μg. Ribosomal RNA depleted, RNase R-treated samples were used as templates for cDNA libraries. RNA-seq libraries were prepared by Illumina Stranded Total RNA Library Prep Kit and then sequenced on the Illumina HiSeq 2000 platform with 2 × 100 bp paired reads. RNA-seq data of human cervical carcinoma HeLa cells were downloaded from the Sequence Reads Archive (accession SRP095410).

### CircRNAs detection, annotation and differential expression analysis

For all RNA-seq data, we followed Song’s computational pipeline UROBORUS (Song et al. 2016) to detect circRNAs. RNA-seq reads were initially mapped to the human reference genome (hg19) using TopHat2 (Kim et al. 2013) (TopHat 2.1.0) with default parameters to filter canonical splicing supporting reads. Unmapped reads of BAM format (“unmapped.bam”) from tophat results were transformed to SAM format (“unmapped.sam”) by SAMtools (Li et al. 2009) (SAMtools 1.3) and then performed detection, annotation as well as statistically analysis of circRNAs from candidate back-spliced junction reads by UROBORUS (UROBORUS 0.1.2 parameter: -index hg19 -gtf hg19_ens.gtf -fasta hg19).

For RNA-seq data of human leukemia samples and cervical carcinoma HeLa cells treated with RNase R, we filtered all back-spliced junction detected by the means number of back-spliced junction reads among all samples and selected those of relative high expression level, and threshold of average one reads per sample was adopted. For RNA-seq data of untreated and *AF10* knock down K562 cells, those back-splicing junctions supported by at least two reads were annotated to be candidate circRNA for each replication in two groups. Differential circRNA expression analysis was performed using the “limma” (Ritchie et al. 2015) and “edgeR” (Robinson et al. 2010) package in R.

### Data access

The CIRCScan assembler is freely available online for academic use. All code and associated data for analysis can be downloaded from GitHub (https://github.com/johnlcd/CIRCScan).

RNA-seq data of human leukemia samples and K562 cells have been submitted to NCBI Sequence Read Archive (SRA) under the accession number of PRJNA397482 (SRA run numbers: SRR6282349-SRR6282354) and PRJNA431686 (SRA run numbers: SRR7475923-SRR7475926). RNA-seq data of human cervical carcinoma HeLa cells were downloaded from GEO database under the accession number of GSE92632.

## Acknowledgements

Not applicable.

## Authors’ contributions

Y. G. and T.-L. Y. designed the study; J.-B. C. and S.-S. D. wrote and revised the manuscript; J.-B. C. performed data analyses; S. Y. and Y.-J. Z. and M.-R. G. tested the pipeline; W.-X. H. and H. C. designed the experiment; Y.-Y. D. and N.-N. W. performed the knocking down experiment; Y.-X. C. and Y. R. collected the GEO data; F.-L. Z. led the AML data sequencing; R.-H. H., H. N. T., Y. G., and T.-L. Y. revised the manuscript.

## Disclosure Declarations

The authors declare that they have no competing interests.

This study was approved by the ethics committee of Xi’an Jiaotong University, and informed consent was obtained from all subjects.

